# Single cell RNA sequencing reveals regulation of fetal ovary development in the monkey (Macaca fascicularis)

**DOI:** 10.1101/2020.05.22.110064

**Authors:** Zheng-Hui Zhao, Chun-Yang Li, Tie-Gang Meng, Yan Wang, Wen-Bo Liu, Ang Li, Yi-Jun Cai, Yi Hou, Heide Schatten, Zhen-Bo Wang, Qing-Yuan Sun, Qiang Sun

## Abstract

Germ cells are vital for reproduction and heredity. However, the mechanisms underlying female germ cell development in primates, especially in late embryonic stages, remain elusive. Here, we performed single-cell RNA sequencing of 12471 cells from whole fetal ovaries, and explored the communications between germ cells and niche cells. We depicted the two waves of oogenesis at single cell resolution and demonstrated that progenitor theca cells exhibit similar characteristics to Leydig cells in fetal monkey ovaries. Notably, we found that *ZGLP1* displays differentially expressed patterns between mouse and monkey, which is not overlapped with *NANOG* in monkey germ cells, suggesting its role in meiosis entry but not in activating oogenic program in primates. Furthermore, the majority of germ cell clusters that highly expressed *PRDM9* and *SPO11* might undergo apoptosis after cyst breakdown, leading to germ cell attrition. Overall, our work provides new insights into the molecular and cellular basis of primate fetal ovary development at single-cell resolution.

## INTRODUCTION

Fetal ovary development is a well-orchestrated complex process that involves the transitions of multiple cell states and communications between germ cells and niche cells. In primates, female germ cell development processes at embryonic stages mainly include oogonia proliferation, meiosis initiation, germ cell attrition and primordial follicle formation, which are accompanied with dynamic chromatin repackaging and transcriptional regulation. Moreover, female germ cells initiate meiosis asynchronously in a wave from anterior to posterior, which results in the heterogeneity of germ cell populations in a fetal ovary ^1^. Somatic progenitor cells in XX gonads will adopt the ovary specific cell fate after sex determination, which leads to their differentiation as granulosa cells in supporting cell populations or as theca cells in steroidogenic cell populations ^2^. Interestingly, the female germ cell fate mainly relies on the ovarian environments established by somatic cells rather than the sex chromosomes in germ cells ^3^.

Despite the mass data produced in recent years on fetal ovary development ^4–6^, several important questions regarding the key developmental events, such as two waves of oogenesis, female germ cell attrition and primordial follicle formation, have not been fully demonstrated, especially in non-human primates. In addition, although the origins of germ cells and granulosa cells are well established ^2, 7^, the origin of theca cells in primates remains unknown. Furthermore, the germline-niche communications through signaling pathways for critical events during fetal ovary development are poorly integrated.

Single cell RNA sequencing approach can efficiently identify cell types, uncover heterogeneity and construct developmental trajectories, which is well suited for exploring fetal ovary development. To further improve our understanding on the germ cell development and somatic cell differentiation, we here performed single-cell RNA sequencing of fetal monkey ovarian cells, especially in late embryonic stages, through 10× Genomics Chromium platform. And a transcriptional cell atlas of all cell types in the fetal ovary was established.

## RESULTS

### Identification of the ovarian cell types using single cell transcriptomes

Single-cell suspensions of fetal Macaca fascicularis ovaries were individually captured and processed with the 10X Chromium system (Figure 1A). Sequencing data from two single cell RNA-seq libraries derived from the two stages of ovary samples were integrated for processing and analysis. From a total of 21036 cells, we obtained an average of 57269 reads per cell and 1593 genes per cell (Figure S1A). Following quality control, 12471 cells were retained for further analysis.

**Figure 1.**
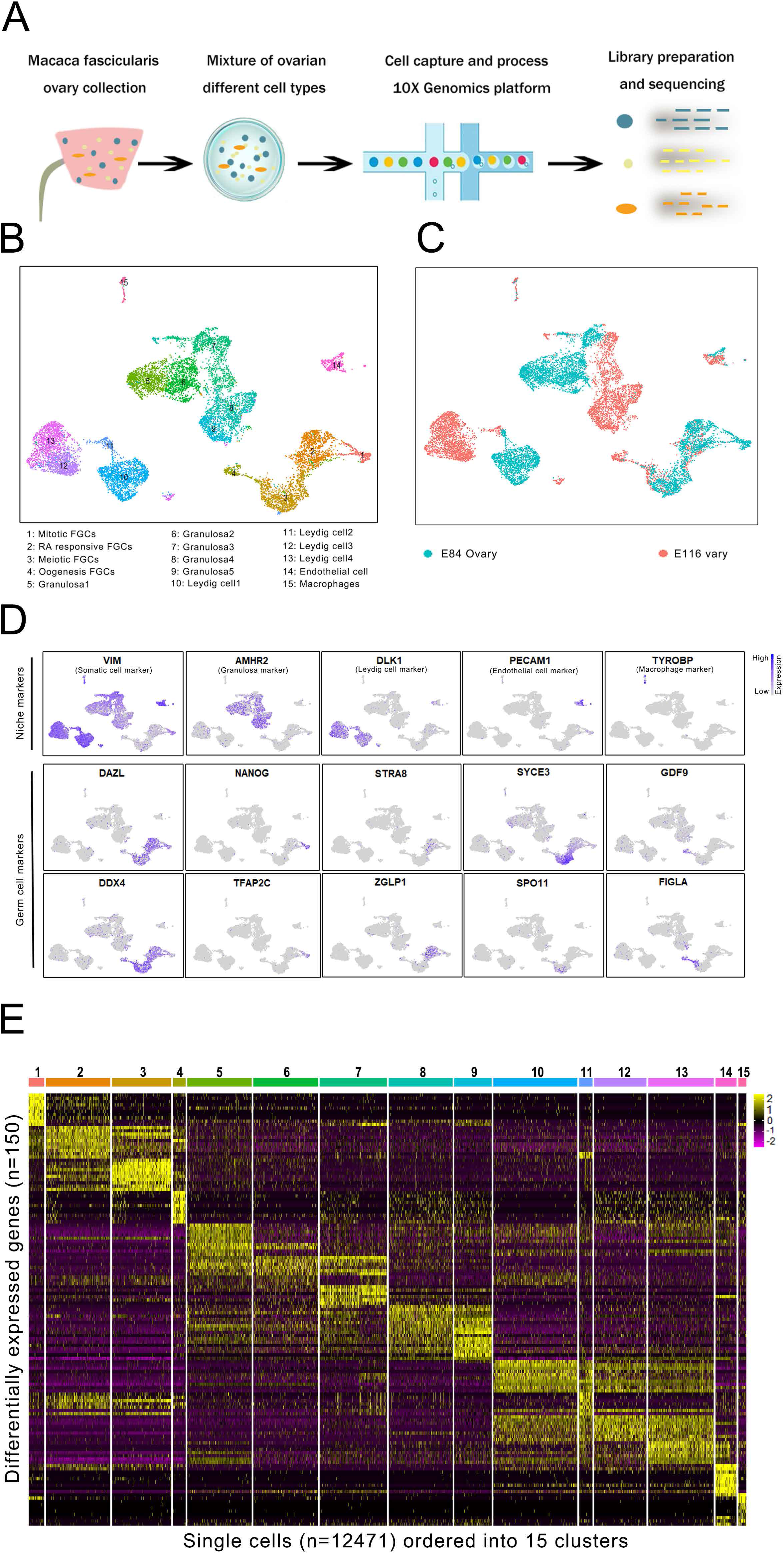
Single cell transcriptome profiling of fetal monkey ovaries. (A) Schematic diagram of the experimental design. (B) Clustering analysis of single cell transcriptome data from fetal monkey ovaries. (C) Single cell transcriptome data with cells colored by embryonic stages. (D) Expression patterns of selected markers for the identification of cell clusters. (E) Heatmap of differentially expressed genes across 15 cell clusters.

To classify the different cell populations in fetal monkey ovaries, we performed unsupervised graph clustering through Seurat package ^8^ to group the cells into cell clusters based on the similarities in their transcriptome profiles. Overall, the 15 transcriptionally distinct clusters were classified and visualized using UMAP (Uniform Manifold Approximation and Projection) (Figure 1B-C) ^9^. To further identify the clusters, we assigned the clusters based on known cell-type marker genes (Figure 1D). The germ cell specific marker genes were expressed solely in clusters 1-4 (*DAZL*, *DDX4*). The clusters 5-15 are somatic cell populations (*VIM*), which include granulosa cells (*AMHR2*), Leydig cells (*DLK1*), endothelial cells (*PECAM1*) and macrophages (*TYROBP*). Also, the germ cell marker genes displayed developmental-stage-specific expression patterns. For instance, primordial germ cell marker gene *TFAP2C* and pluripotency marker gene *NANOG* were expressed in the early mitotic female germ cells (FGCs), whereas *STRA8* was expressed in retinoid acid (RA) signaling-responsive FGCs. The cells in cluster 3 were meiotic prophase FGCs and specifically expressed *SYCE3* and *SPO11*. Finally, the cells in cluster 4 were oogenesis phase FGCs and clearly expressed *GDF9* and *FIGLA*. Collectively, the sequential expressed germ cell markers in clusters 1 to 4, respectively could mirror the temporal order of early female germ cell development. Interestingly, our results also demonstrated multiple transcriptionally distinct sub-populations within the granulosa cells (five clusters) and Leydig cells (four clusters). Additionally, the top 10 differentially expressed genes were selected across cell clusters for the identification of each cell type (Figure 1E). Together, we have profiled the transcriptomes of all cell types in the fetal monkey ovary in the present study.

### Cell lineage reconstruction reveals two waves of oogenesis

To explore the female germ cell development at a higher resolution, we isolated and re-clustered the germ cell populations, which were visualized using UMAP (Figure 2A-B). To further recapitulate the trajectory of early female germ cell development, we constructed the germ cell lineage using Monocle2 package ^10^, which organized the cells along the pseudotime trajectory, with mitotic FGCs and oogenesis phase FGCs concentrated at the beginning and end of its axis, respectively (Figure 2C).

**Figure 2.**
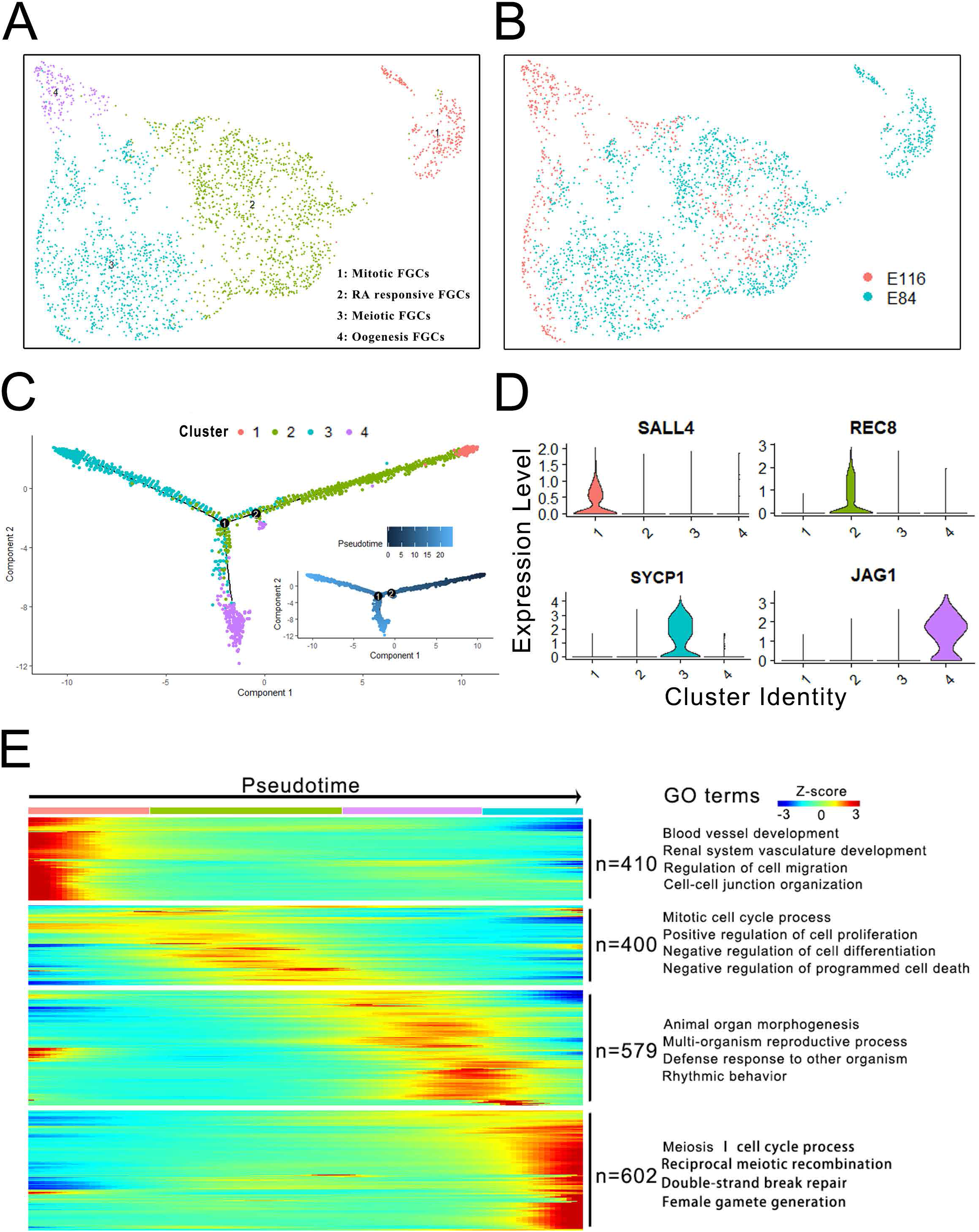
Cell lineage reconstruction and identification of the germ cells. (A) Re-clustering of the germ cell populations. (B) Single cell RNA sequencing data with germ cells colored by embryonic stages. (C) Pseudotime analysis of germ cells. (D) Violin plots show the expression patterns of marker genes during oogenesis. (E) Hierarchical clustering of the ordering genes during early oogenesis.

The reconstruction of the germ cell lineage divided the germ cell lineage into three developmental windows defined by the branch points of the pseudotime trajectories (Figure 2C), which allowed us to identify transition states leading to the oogenesis. We found that two waves of oogenesis occur in the germ cell lineage at the branch points 1 and 2. Noticeably, the first wave of oogenesis occurs following the RA responsive stage, whereas the second wave of oogenesis occurs in the late meiotic prophase (Figure 2C). The first wave of oogenesis may contribute to the medullary follicles that are activated immediately after their assembly, while the second wave of oogenesis contributes to the cortical primordial follicles that are activated after puberty and recruited regularly under hormonal control ^11, 12^. To further determine the patterns of differentially expressed genes, we examined the stage-specific marker genes, such as *SALL4*, *REC8*, *SYCP1*/*2* and *JAG1*, which change dynamically over the trajectory and are involved in oogonia proliferation, meiosis initiation, meiosis progression and primordial follicle formation, respectively (Figure 2D, Figure S1B). Also, to dissect the features of various stages of early female germ cell development, 2000 ordering genes that expressed dynamically along pseudotime were selected and clustered. The heatmap revealed stage-specific gene expression patterns that were consistent with well-organized germline development (Figure 2E). As expected, gene ontology (GO) analysis of the clustered ordering genes identified several significantly enriched biological processes, such as “mitotic cell cycle process” and “meiosis | cell cycle process”. In summary, these data displayed the dramatic changes of transcriptomes during early germ cell development.

### Dynamic transcription factors control meiosis initiation and progression

Having identified the cell clusters and constructed the developmental trajectory of FGCs, we next determined the transcriptome of FGCs during their development. To explore the transition from mitosis to meiosis, we clustered the mitotic and meiotic genes across germ cell populations. As expected, mitotic genes, such as *RCC2* and *MYBL2*, mainly expressed in cluster 1, while meiotic genes, such as *RAD17*, *TEX11* and *MEIOB*, mainly expressed in cluster 3 (Figure 3A). Also, the *CENPF*, *PRKDC* and *CHMP2A* exhibited peak expression in RA responsive FGCs, which may play important roles in meiosis entry. Additionally, mitotic phase FGCs clearly express pluripotency markers such as *NANOG* and *LIN28A* ^13, 14^ (Figures S1B-C). Subsequently, the total expression levels of pluripotent genes significantly decreased during the transition from mitosis to meiosis, while meiotic marker genes, such as *STRA8*, *DMC1* and *INCA1*, increased strikingly (Figure 3B, Figure S1C). In oogenesis phase FGCs, maternal effect genes, such as *FIGLA*, *ZAR1* and *ASTL*, begin to express (Figure 3B, Figure S1C), suggesting the start of primordial follicle formation and folliculogenesis.

**Figure 3.**
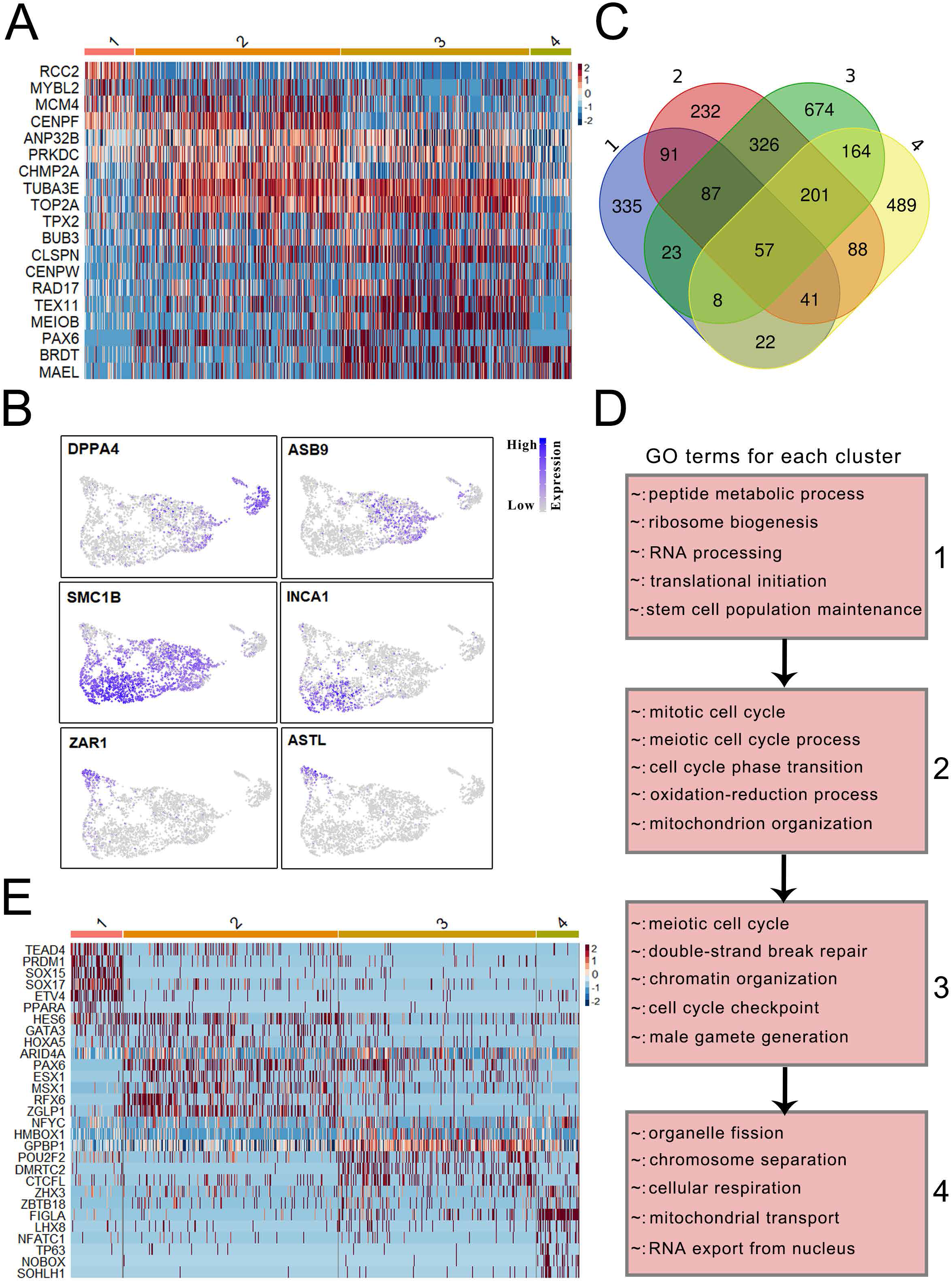
Mitosis to meiosis transition and meiosis progression. (A) Heatmap of differentially expressed mitotic and meiotic genes. (B) Expression patterns of germ cell development associated genes. (C) Overlap of differentially expressed genes across germ cell clusters. (D) Differentially expressed genes and associated GO categories characteristic of meiosis initiation and progression. (E) Heatmap representing the transcription factors across germ cell clusters.

To further dissect the differences in gene expression profiles, we detected 664, 1123, 1540, and 1070 differentially expressed genes among the first four clusters, respectively. And these differentially expressed genes changed significantly during the mitosis to meiosis transition and meiosis to oogenesis transition (Figure 1E, Figure 3C). Also, we performed GO analysis on the differentially expressed genes in each of the germ cell clusters. The genes in cluster 1 were enriched in the GO terms of “peptide metabolic process” and “stem cell population maintenance”, while the genes in cluster 3 were enriched in the categories of “meiotic cell cycle” and “chromatin organization involved in meiosis”, which showed striking meiosis-related features (Figure 3D). Noticeably, RA responsive phase FGC-specific genes were enriched in mitotic and meiotic cell cycle processes, suggesting that the cell cycle phase transition occurs in this stage.

To further explore the regulation of meiosis initiation and progression, we analyzed the differentially expressed transcription factors across germ cell clusters (Figure 3E). We found that transcription factor genes like *TEAD4*, *SOX4*, *ETV4*, *PRDM1*, *SOX15* and *SOX17* showed expression peaks specifically in the mitotic stage FGCs, which may be involved in primordial germ cell proliferation. Additionally, *PAX6*, *ESX1*, *MSX1*, *RFX6* and *ZGLP1* exhibited peak expression in RA responsive stage FGCs. Evidence has shown that *MSX1* and *ZGLP1* are associated with meiosis initiation ^15, 16^, but no reports exist yet about the functions of *PAX6*, *ESX1* and *RFX6* in meiosis. Noticeably, *DMRTC2* specifically expressed in cluster 3 (Figure 3E, Figure S1D), which might play an important role in meiosis progression ^17^. For cluster 4, cells showed high levels of folliculogenesis-associated genes such as *FIGLA*, *NOBOX* and *SOHLH1*. Furthermore, the zinc finger protein family plays important roles in early oogenesis. For example, *ZNF625*, *ZNF217* and *ZNF728* mainly expressed in cluster 1, while *ZNF131* and *ZNF711* exhibited peak expression in cluster 3. Overall, our data demonstrate that the different germ cell clusters may represent different cell stages en route to early oogenesis.

### Majority of female germ cells adopt an apoptotic fate

In contrast to the second wave of oogenesis, the majority of female germ cells undergo apoptosis after the branch point 1 (Figure 2C). To further explore the mechanisms of germ cell fate determination, we performed re-clustering of the germ cells at a higher resolution and reconstructed the germ cell lineage using Monocle2. As a result, 11 sub-clusters were identified and the G8 and G9 may likely undergo apoptosis, while the G7 and G10 may generate primordial follicles (Figure 4A-B). Also, the G8 and G9 at the end of the pseudotime exhibited sharply increased expression of meiosis-associated genes, such as *SYCP1* and *HORMAD1*, while G7 and G10 highly expressed *MT1X* that is essential for anti-apoptosis (Figure 4C, Figure S2A) ^18^. To identify genes that exclusively expressed in a given cell type, we performed differential gene expression analysis to identify highly variable genes for each cluster. Venn diagram shows the significant differences between G7/G10 and G8/G9 (Figure 4D).

**Figure 4.**
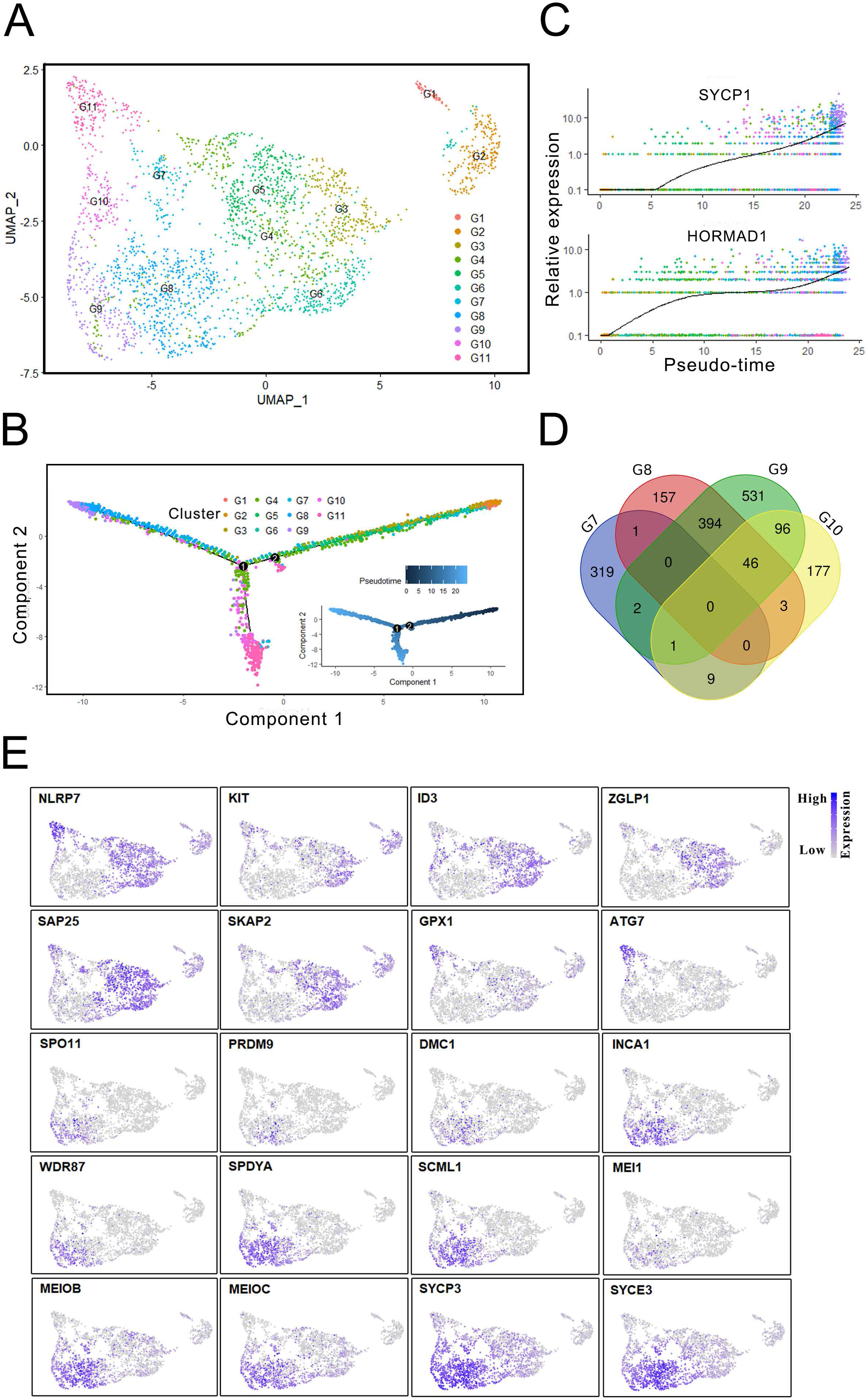
Female germ cell fate determination. (A) Re-clustering of germ cells at a higher resolution. (B) Cell lineage reconstruction through monocle. (C) Dynamic expression of meiotic genes. (D) Overlap of highly variable genes among G7 to G10 clusters. (E) The expression patterns of germ cell fate determination-associated genes.

To further dissect the differences between apoptotic and oogenic germ cells, we selected several marker genes that differentially expressed among these clusters (Figure 4E). For example, *NLRP7* is specifically expressed in germ cells that undergo oogenesis ^19^, but this gene is not expressed in G8 and G9, which indicated that the cells in cluster G8 and G9 were probably primed for the apoptosis process. Moreover, several genes associated with germ cell development, such as *KIT* that is essential for germ cell survival ^20^, also displayed similar expression patterns with *NLRP7*. Furthermore, *ZGLP1* was shown to induce the oogenic fate in mice^16^, which is absent in cluster G8 and G9 but expressed in other meiotic germ cells. In contrast, meiotic genes, such as *SPO11*, *PRDM9* and *DMC1*, specifically expressed in G8 and G9 clusters. And PRDM9-mediated H3K4me3 at hotspots could direct double-strand breaks (DSBs) fate during meiosis recombination ^21^. Additionally, *WDR87*, *SPDYA* and *SCML1* also exhibited higher expression in G8 and G9, which may be critical for the regulation of germ cell fates (Figure 4E). Collectively, our findings indicate that the majority of FGCs are probably ready to undergo apoptosis, which may contribute to female germ cell attrition ^22^.

### Identification of the granulosa cell subpopulations

Granulosa cells are the most important somatic cells in the ovary, which markedly express *AMHR2*, *FOXL2* and *FST* genes (Figure 5A), and play important roles in female germ cell development and in the formation of primordial follicles. To examine the features of granulosa cells more closely, we detected the differentially expressed genes in each cluster, and the highly variable expressed genes in granulosa cells were identified (Figure 5B). For example, *HES1*, *NR4A1* and *EGR3* mainly expressed in cluster 5, while *ADIRF*, *CBLN2* and *LYPD1* exhibited peak expression in cluster 9. Next, we performed principal component analysis (PCA) on the significant principal components to compare the transcriptomes of granulosa cells (Figure S2B). The PC1 axis had the most differences between the two samples. The E84 sample was located on the left side, whereas the E116 sample was located on the right side. The PC2 axis was dominated by differences in the cluster level within granulosa cells. However, we did not find any distinct boundary to group the granulosa cells.

**Figure 5.**
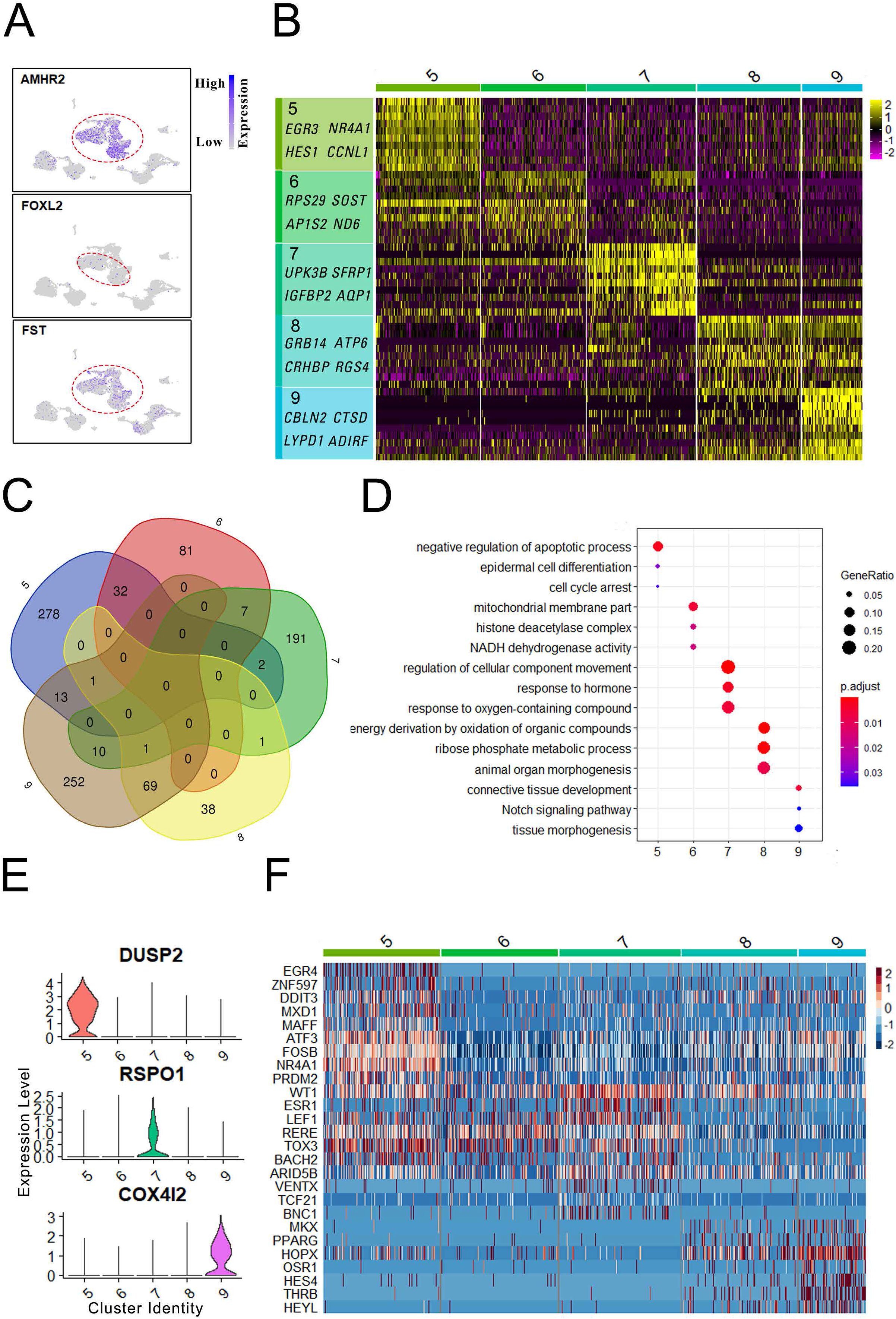
Granulosa cell transcriptome dynamics. (A) Feature plots of granulosa cell marker genes. (B) Expression patterns of differentially expressed genes across granulosa cell populations. (C) Venn diagram shows overlapping of differentially expressed genes among granulosa cell populations. (D) The enriched biological processes to the subpopulations of the granulosa cells. (E) Violin plots show the highly variable genes among granulosa cell subpopulations. (F) Heatmap representing the dynamics of transcription factors among granulosa cell populations.

To further explore the differences among granulosa cell clusters, we detected 326, 120, 212, 110 and 346 genes that differentially expressed across granulosa cell populations. Venn diagram shows that there are significant differences between cluster 5 and cluster 8 as well as cluster 6 and cluster 9 (Figure 5C). This result suggests that the granulosa cells have undergone dramatic changes from E84 to E116 stages. To ascertain the features of granulosa cell subpopulations, we performed GO analysis on the differentially expressed genes in each cluster (Figure 5D). As a result, the genes in cluster 5 were enriched in the categories of “negative regulation of apoptotic process” and “cell cycle arrest”, while the genes in cluster 9 were enriched in the GO terms of “notch signaling pathway” and “tissue morphogenesis”. Noticeably, certain genes in cluster 7 were enriched in the categories of “response to hormone”. Collectively, these results indicated that subpopulations of granulosa cells play distinct roles in fetal ovary development.

To dissect the mechanisms that regulate granulosa cell development, we analyzed the differentially expressed genes and transcription factors across granulosa cell clusters. We found that highly variable genes like *RSPO1*, *DUSP2* and *COX4I2* exhibited peak expression specifically in the cluster 5, cluster 7 and cluster 9, respectively (Figure 5E). Noticeably, *RSPO1*, an activator of the WNT/β-catenin pathway, is located upstream of the female sex determination signaling pathway, which could promote cell proliferation and ovarian differentiation ^23, 24^. In contrast, *DUSP2* is a substrate for mitogen-activated protein kinases (MAPKs) ^25^, which may play an important role in inhibiting cell proliferation. Additionally, *COX4I2* is specifically expressed in cluster 9, which is involved in the modulation of oxygen affinity through hypoxia-sensing pathways, suggesting its role in the activation of primordial follicles ^26^. Interestingly, *RDH10* also expressed in cluster 9 (Figure S2C), which is essential for the production of RA ^27^. On the other hand, transcription factors were clustered across granulosa cell subpopulations (Figure 5F). The *EGR4*, *ZNF597*, *DDIT3*, *MXD1* and *MAFF* specifically expressed in cluster 5, while *MKK*, *PPARG*, *OSR1*, *HES4*, *THRB* and *HEYL* mainly expressed in cluster 8 and cluster 9. Additionally, *ATF3*, *FOSB* and *PRDM2* displayed higher expression in cluster 5 than other populations, whereas *ESR1* exhibited peak expression in cluster 7, which is consistent with the “response to hormone” category in cluster 7. Collectively, highly variable expressed transcription factors may instruct the differentiation of granulosa cells in a distinct manner.

### Similar characteristics between progenitor theca cells and fetal Leydig cells

Theca cells are another important somatic cell type in the ovary, which is required for folliculogenesis. However, the origin of theca cells has not been fully demonstrated. To explore the origin of theca cells, we performed single cell RNA sequencing on the whole fetal ovary. Surprisingly, in addition to granulosa cells, there are four groups of cells that highly expressed fetal Leydig cell markers, such as *DLK1*, *CXCL12*, *PDGFRA* and *TCF21* (Figure 6A). Therefore, we named these cells Leydig cells. Noticeably, Leydig cells also express the genes *NR2F2*, *WT1* and *GLI1* (Figure 6A), which suggested that Leydig cells may be the progenitor theca cells ^2, 28^. To further determine the origin of theca cells, we re-clustered the Leydig cells at a higher resolution and detected the levels of theca cell marker genes. As expected, the structural theca cell marker genes *PTCH1* and *ACTA2* as well as endocrine theca cell marker gene *CYP17A1* exhibited differentially expressed patterns across Leydig cell populations (Figure 6B), which indicated that progenitor theca cells exhibit the similar characteristics to fetal Leydig cells.

**Figure 6.**
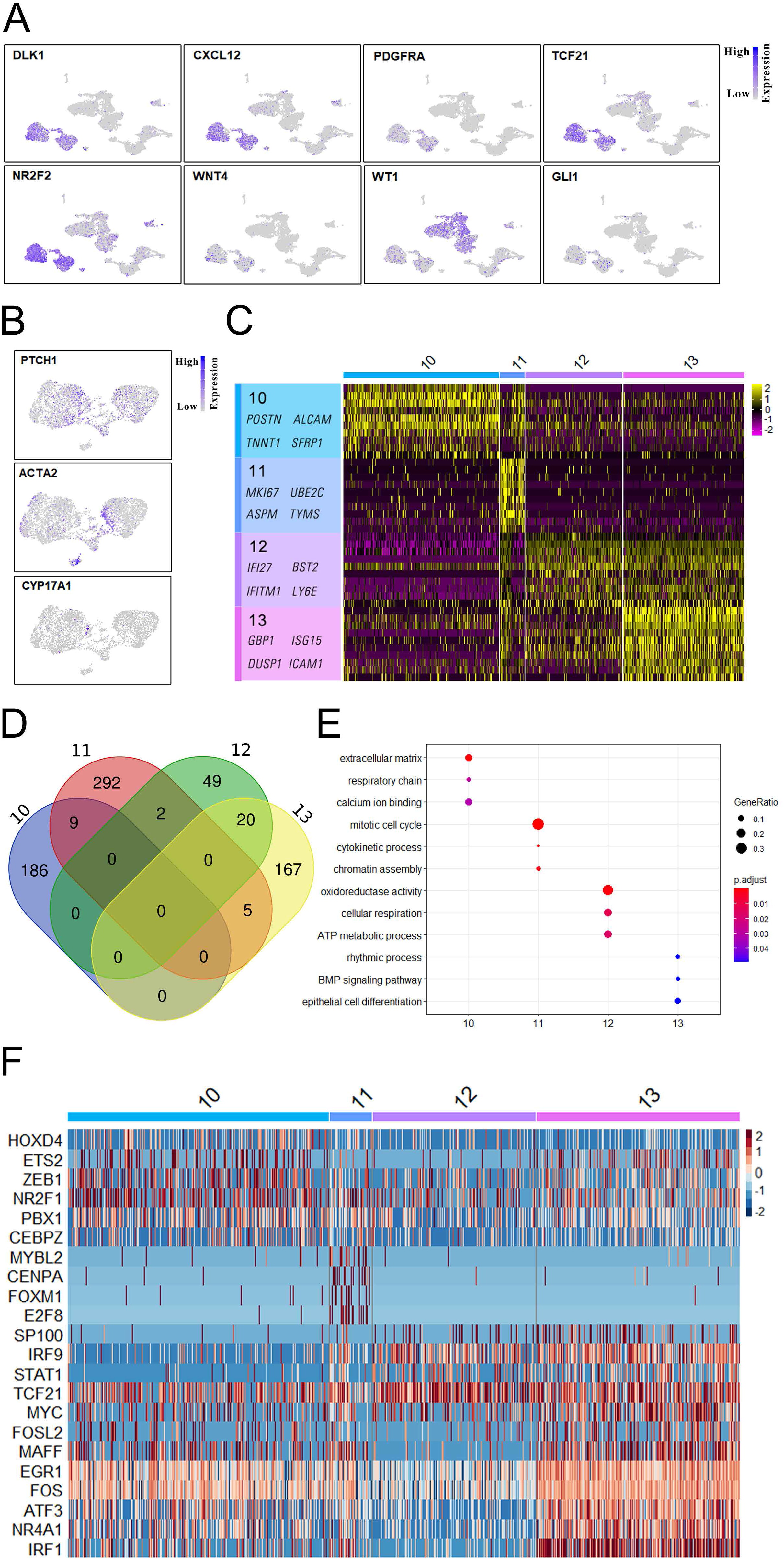
Dynamic transcriptome in theca cells. (A) Expression profiles of known marker genes for Leydig cells. (B) The expression patterns of theca cell-associated marker genes. (C) Expression patterns of differentially expressed genes across theca cell populations. (D) Venn diagram shows overlapping of differentially expressed genes among theca cell populations. (E) Top GO terms within the differentially expressed genes unique to the theca cells. (F) Heatmap representing the dynamics of transcription factors among theca cell subpopulations.

To further characterize theca cells, we detected the highly variable expressed genes across theca cell populations (Figure 6C; Figure S3A). Obviously, cluster 11 is significantly different from the other three clusters. For example, *MKI67*, *SGO1*, *CDK1*, *CCNA2* and *CENPF* specifically expressed in cluster 11, whereas *IFI27*, *IFITM1* and *BST2* generally expressed in cluster 12 and cluster 13 (Figure 6C; Figure S3A). Additionally, 195, 308, 71 and 192 differentially expressed genes were detected across theca cell populations. Similar to granulosa cells, theca cells exhibited distinct features between each cluster (Figure 6D). To explore the features of theca cells, we carried out GO analysis on the differentially expressed genes in each cluster (Figure 5E). Noticeably, the genes in cluster 13 were enriched in GO terms of “rhythmic process” and “BMP singling pathway”, which suggests that theca cells in cluster 13 may be involved in folliculogenesis. To dissect the mechanisms that control theca cell differentiation, we clustered the transcription factors across theca cell populations. Obviously, the transcription factors in cluster 11, such as *MYBL2*, *CENPA*, *FOXM1* and *E2F8*, displayed significant differences compared to that in the other three theca cell clusters (Figure 6F). The fate of cells in cluster 11 need to be further explored.

### Signaling pathways and interactions between niche and germ cells throughout fetal ovary development

The mitosis to meiosis transition and primordial follicle formation processes are crucial steps during fetal ovary development in primates. However, the molecular mechanisms and signaling pathways that are involved in these processes have not been fully explored. Therefore, we investigated the changes in RNAs encoding signaling pathwayassociated factors during fetal ovary development, to provide insights into female germ cell development and niche-germ cell interactions.

Our sequencing data showed that the RA signaling pathway and bone morphogenic protein (BMP) signaling pathway are involved in oogenic program initiation and meiotic progression. For the BMP signaling pathway, the ligand *BMP2* was highly expressed in granulosa cells, whereas the receptor *BMPR1B* was expressed in both granulosa cells and FGCs. Noticeably, the targets *ID1*, *ID2* and *ID3* were expressed in FGCs in a stage-specific manner (Figure 7A). This pattern suggests that the BMP signaling pathway may play an important role in meiosis initiation and progression. Additionally, the ligand of the KIT signaling pathway (*KITLG*) was specifically expressed in granulosa cells, and the receptor *KIT* was highly expressed in FGCs, which is crucial for germ cell survival ^20^. Interestingly, *PDGFB* was specifically expressed in endothelial cells, and its receptors *PDGFRA* and *PDGFRB* were found in theca cells and partial granulosa cells (Figure S3B), suggesting that endothelial cells may indirectly affect female germ cell development, through the interactions with other niche cells.

**Figure 7.**
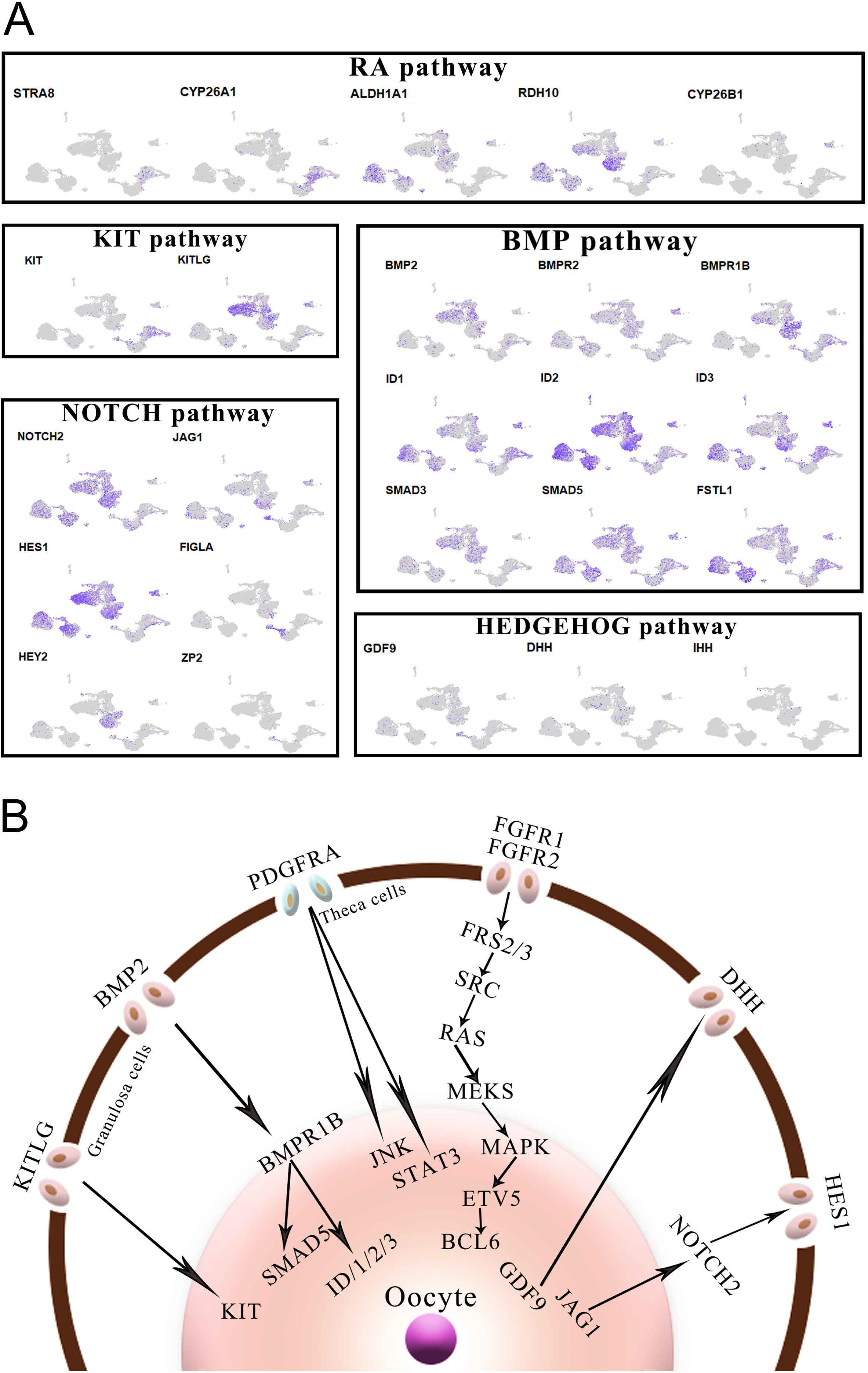
Signaling pathways for niche-germline interactions. (A) Relative expression levels of marker genes from different key signaling pathways. (B) Schematic summary of signaling pathways for niche-germline interactions.

To further explore the interactions between the oocyte and granulosa cells throughout folliculogenesis, we analyzed the expression of the factors in key signaling pathways, including NOTCH and HEDGEHOG pathways. Our results showed that the ligand *JAG1* of NOTCH pathway was predominantly expressed in the oocytes at the oogenesis phase, whereas the receptor *NOTCH2* as well as the downstream target gene *HES1* were highly expressed in granulosa cells and progenitor theca cells (Figure 7A). Next, we investigated the key components of the HEDGEHOG signaling pathway in the oocytes and somatic cells. *GDF9* exhibited high expression level in the oogenesis phase FGCs, which could promote the production of Desert hedgehog (DHH) and Indian hedgehog (IHH) in granulosa cells. Moreover, the receptor *PTCH1* as well as downstream signaling components *GLI1* of the HEDGEHOG pathway were highly expressed in theca cells (Figure 6A-B), indicating that the HEDGEHOG pathway may play critical roles in the specification of theca cells. Taken together, our results demonstrate that reciprocally interacting signaling pathways control fetal ovary development in a stage-specific manner.

## DISCUSSION

Fetal ovary development is a complex process in primates, and a full understanding of its regulation requires the integration of multiple data collected from various cell types in the ovary. Here, we performed single cell RNA sequencing analysis of all cells within the fetal ovaries at two different developmental stages to provide new insights into the regulation of fetal ovary development in primates. In this study, we have identified four clusters of female germ cells: mitotic phase, RA responsive phase, meiotic phase and oogenesis phase. Moreover, the niche cells, such as granulosa cells, theca cells, endothelial cells and macrophages, were also identified. Furthermore, our work revealed unique characteristics in transcriptional profiles and reciprocal interactions between germ cells and niche cells in each developmental stage. These identified cell types and stage-specific expressed genes in fetal ovaries may offer a valuable information for future functional studies.

Female germ cells undergo meiosis initiation, primordial follicle formation and apoptosis. Here, we reconstructed the developmental trajectory of female germ cells, which revealed two waves of oogenesis. Noticeably, the first minor oogenesis occurs following the RA responsive stage, which could further undergo growth and development in the medulla ^12^. However, these follicles have to undergo atresia due to lacking hormone stimulation in the prepuberty ovary, and cannot contribute to fertility. On the other hand, the second major oogenesis occurs in the late meiotic prophase accompanied with the apoptosis of the vast majority of germ cells. The germ cells in the second oogenesis will form primordial follicles and stay dormant in the ovarian cortex until puberty ^11^. After puberty, the primordial follicles are recruited regularly to generate mature oocytes under the stimulation of gonadotropin hormones, which contributes to the life span of female reproduction.

The mitosis to meiosis transition is a critical step in the oogenic program, however, the mechanisms for the regulation of meiosis initiation have not been fully demonstrated. Previous studies have shown that RA induces the expression of *STRA8*, which could further switch on the meiotic genes, such as *SYCP3*, and repress the early PGC program to promote meiosis initiation. RA enhances this pathway through retinoic acid receptor (RAR) -mediated transcriptional regulation; however, after ablation of RAR-coding genes, the female germ cells robustly expressed meiotic genes, such as *STRA8*, *REC8* and *SYCP3*, showing that RARs are actually dispensable for meiosis initiation ^29^. Moreover, recent studies indicate that *Zglp1* is partially overlapped with *Nanog* in mice, which is sufficient to induce the oogenic fate and meiosis entry, whereas RA augments the ZGLP1-activated oogenic program and meiosis initiation^16, 30^. In our study, *ZGLP1* mainly expressed in germ cells at the RA responsive stage, which is not overlapped with *NANOG* in monkey female germ cells, indicating that *ZGLP1* is essential for meiosis entry but not for activating the oogenic fate in primates. Moreover, *ZGLP1* also expressed in the oogenesis phase germ cells, suggesting its roles in subsequent oogenesis.

During the second wave of oogenesis the germ cells are surrounded by somatic cells to form primordial follicles, while the vast majority of germ cells undergo apoptosis. Reconstructed germ cell lineage revealed that C8 and C9 are likely to undergo apoptosis, while other germ cells maintain survival and undergo oogenesis. *NLRP7* generally expressed in all tstages of oogenesis, and *KIT* is crucial for the survival of germ cells ^19, 20^. However, *NLRP7* and *KIT* did not express in the C8 and C9. Noticeably, several meiosis-associated genes, such as *SPO11*, *PRDM9*, *DMC1* and *INCA1*, exhibited peak expression in C8 and C9. Recent studies have shown that PRDM9-mediated H3K4me3 could guide SPO11 targeting to induce DSBs ^31^. Moreover, earlier formed DSBs occupy more open chromatin and are much more competent to proceed to a crossover fate, whereas later formed DSBs are likely to proceed to a non-crossover fate ^21^. In this study, *PRDM9* and *SPO11* specifically expressed in C8 and C9, whose DSBs are formed late causing their non-crossovers fate, which may be the cause for apoptosis of germ cells in C8 and C9.

Ovary and testis have the same developmental origin: the bipotential gonads that are composed of multipotent somatic progenitor cells. After sex determination, the somatic progenitor cells differentiate into Sertoli cells and Leydig cells in male gonads, or granulosa cells and theca cells in female gonads. Our data revealed five clusters of granulosa cells that displayed distinct features in late stage ovaries. In addition, we also provide several new insights into theca cell development, most importantly into the origin of theca cells. Apart from the somatic progenitor cells that differentiated into granulosa cells, the remaining somatic progenitor cells exhibited peak expression of *NR2F2*, which acquire a steroidogenic precursor fate by progressively expressing *CYP17A1*, *PTCH1* and *ACTA2*. Interestingly, these cells also exhibited the Leydig cell features, which highly expressed Leydig cell marker genes, such as *DLK1*, *TCF21* and *CXCL12*. In contrast to *SRY* that is essential for male sex determination, *NR2F2* has been shown to control female sex determination through eliminating Wolffian ducts in female embryos ^32^. Moreover, *NR2F2*, a lineage-specific transcription factor, plays important roles in cell type specification and cell fate maintenance ^33^. Therefore, we speculated that progenitor theca cells exhibit similar features to fetal Leydig cells, and the features of Leydig cells are eliminated gradually through *NR2F2* with the development of the ovary.

A major area of current interest involves niche-germline communications that coordinately and reciprocally regulate fetal ovary development, but knowledge on how the signaling pathways interact remains limited. In this study, we have identified many ligands and receptors that are derived from the reciprocal compartments. For example, we found that the BMP signaling pathway was activated in female germ cells via granulosa cells driven mechanisms. BMP2, the ligands for the BMP signaling pathway, specifically expressed in granulosa cells, while its receptor, BMPR1B, and downstream effector, *ZGLP1*, expressed in female germ cells. Our data concur that the BMP-ZGLP1 pathway could also activate the oogenic program in primates. Moreover, typical RA pathway contributes to oogenic program maturation and PGC program repression. In contrast, NOTCH and HEDGEHOG pathways were activated in niche cells. For instance, the ligand *JAG1* of NOTCH pathway specifically expressed in the oocytes, while its receptor *NOTCH2* as well as the downstream target gene *HES1* were highly expressed in granulosa cells and progenitor theca cells. In this study, we have identified several pathways that may govern the fetal ovary development in primates, which provides valuable information for the improvement of in vitro culture of gametes.

In summary, this work provides new insights into the crucial features of monkey fetal ovaries especially in the late embryonic stages. Our study paves the way for understanding the molecular regulation of fetal female germ cell development, including meiosis initiation, primordial follicle formation and apoptosis of germ cells. It also lays a solid foundation for the theca cell origin identity. More importantly, the reciprocal relationship between the signaling pathways of FGCs and their niche cells will provide a valuable resource for further optimizing and improving the efficiency of germ cell differentiation in vitro.

## MATERIALS AND METHODS

### Animal Ethics Statements

Fetal female cynomolgus monkeys (Macaca fascicularis) were selected for this study. The use and care of animals complied with the guideline of the Animal Advisory Committee at the Institute of Neuroscience, Chinese Academy of Sciences. The ethics application entitled ‘‘Construction of cynomolgus monkey model based on somatic cell nuclear transfer’’ (#ION-2018002R01) was approved by the Institute of Neuroscience, Chinese Academy of Sciences.

### Sample collection

The ovary samples for scRNA-seq were from two fetal female Macaca fascicularis (embryonic day 84 and 116). For each single cell sequencing experiment, two ovaries were washed twice in 1× PBS, and subjected to a standard digestion procedure through Tumor Dissociation Kit, human (Miltenyi Biotec # 130-095-929). First, the ovaries were cut into small pieces of 2–4 mm, and transferred into the gentleMACS C Tube containing the enzyme mix (4.7 mL DMEM, 200 μL Enzyme H, 100 μL Enzyme R, and 25 μL Enzyme A). Then, the C Tube was attached onto the sleeve of the gentleMACS dissociator. After termination of the program, the C Tube was detached from the gentleMACS Dissociator. The sample was incubated for 30 minutes at 37 °C under continuous rotation using the MACSmix Tube Rotator. Next, the C Tube was attached onto the sleeve of the gentleMACS dissociator, and then subjected to a short centrifugation step to collect the sample material at the bottom of the tube. The sample was resuspended and the cell suspension was applied to a MACS SmartStrainer (70 μm) placed on a 50 mL tube and washed in a cell MACS SmartStrainer (70 μm) with 20 mL DMEM. Finally, the cell suspension was centrifuged at 300×g for 7 minutes, the supernatant was completely aspirated and cells were resuspended as required for further applications.

### Construction of single-cell RNA libraries and sequencing

Single-cell suspensions of ovary cells were captured on a 10X Chromium system (10X Genomics). Then, single-cell mRNA libraries were generated using the single-cell 3’ reagent V3 kits according to the manufacturer’s protocol. About 16000 cells were added to each channel with a targeted cell recovery estimate of 8000 cells (10000 for E84 monkey and 12947 for E116 monkey). After generation of GEMs, reverse transcription reactions were barcoded using a unique molecular identifier (UMI) and cDNA libraries were then amplified by PCR with appropriate cycles. Subsequently, the amplified cDNA libraries were fragmented and then sequenced on an Illumina NovaSeq 6000 (Illumina, San Diego).

### Single cell RNA-seq data processing and analysis

The Cell Ranger software (version 3.0.2) was used to perform sample demultiplexing, reads mapping and barcode processing to generate a matrix of gene counts versus cells. Briefly, the raw BCL files generated by Illumina NovaSeq 6000 sequencing were demultiplexed into fastq files through the Cell Ranger *mkfastq* pipeline. Next, the fastq files were processed using the Cell Ranger *count* pipeline to map high-quality reads to the monkey reference genome (Macaca_fascicularis_5.0). Aligned reads were further filtered for valid cell barcodes and unique molecular identifiers (UMIs) to produce a count matrix.

Then, count matrix was imported into the R package Seurat ^8^ and quality control was performed to remove outlier cells and genes. Cells with 200-3000 detected genes were retained. Genes were retained in the data if they were expressed in ≥ 3 cells. After applying these quality control criteria, 11742 cells and 19204 genes remained for downstream analyses. Additional normalization was performed in Seurat on the filtered matrix to obtain the normalized count. Highly variable genes across single cells were identified and principal component analysis (PCA) was performed to reduce the dimensionality on the top 18 principal components. Then, cells were clustered at a resolution of 0.6 and visualized in 2-dimensions using Uniform Manifold Approximation and Projection (UMAP) ^9^.

### Pseudotime analysis of single-cell transcriptomes

Germ cell lineage trajectories were constructed according to the procedure recommended in the Monocle2 documentation (http://cole-trapnell-lab.github.io/monocle-release/docs) ^10^. Cluster 1 (mitotic PGCs) was used as start, cluster 2 (meiotic PGCs) as middle, and cluster 4 (early activated oocytes) as the end of pseudotime. Briefly, the top differentially expressed genes were selected as ‘ordering genes’ to recover lineage trajectories in Monocle2 using default parameters. After pseudotime time was determined, differentially expressed genes were clustered to verify the fidelity of lineage trajectories. Additionally, the expression patterns of key germ cell markers across pseudospace were visualized through the function of plot_genes_in_pseudotime in Monocle2.

### Gene ontology analysis

Enrichment analysis for highly variable genes detected per cluster was conducted using ClusterProfiler R package ^34^. Symbol gene IDs were translated into Entre IDs through bitr function. The analysis of the enrichment of differentially expressed genes was performed and a corrected P-value≤ 0.05 was considered to indicate significant gene enrichment.

## DATA AVAILABILITY

The high-throughput sequencing data in this study have been deposited in the Gene Expression Omnibus (GEO) database under accession number GSE149629.

To review GEO accession GSE149629:

Go to https://www.ncbi.nlm.nih.gov/geo/query/acc.cgi?acc=GSE149629

Enter token exkjqgaibhyfhqt into the box.

## ACKNOWLEDGEMENTS

We appreciate help from members of the Sun laboratory for comments during preparation of the manuscript. We thank Fei-Yang Wang from State Key Laboratory of Stem Cell and Reproductive Biology, Institute of Zoology, Chinese Academy of Sciences for assistance with bioinformatic analysis.

## AUTHOR CONTRIBUTIONS

Q.-Y.S. and Q.S. conceived the project and designed the experiments. Z.-H.Z. conducted the experiment and performed the data analysis. Sample collection was led by C.-Y.L and Y.W.. T.-G.M., W.-B.L., A.L., Y.-J.C. and Y.H. provided the technical support. Q.-Y.S. and Z.-H.Z. wrote the paper. Q.-Y.S., H.S. and Z.-B.W. edited the manuscript. All authors read and approved the final manuscript.

---

**Figure S1. Sequencing information and differentially expressed genes in germ cell clusters. (A) Sequencing information including cell number, mean reads per cell and median genes per cell etc. (B) Violin plots show the expression patterns of marker genes across germ cell clusters. (C) Feature plots of the stage-specific expressed genes in germ cell clusters. (D) Violin plots of the critical transcription factors across germ cell populations. (E) Heatmap representing the zinc finger protein family genes across germ cell clusters.**

**Figure S2. Highly variable genes in germ cell and somatic cell clusters. (A) Dynamic expression of highly variable genes along pseudo-time. (B) The expression patterns of highly variable genes among granulosa cell subpopulations. (C) PCA analysis of the transcriptome of granulosa cells.**

**Figure S3. Differentially expressed genes and signaling pathways. (A) Highly variable genes among theca cell subpopulations. (B) The expression patterns of marker genes from different key signaling pathways.**

## REFERENCES

1 Bullejos M, Koopman P. Germ cells enter meiosis in a rostro-caudal wave during development of the mouse ovary. Molecular reproduction and development 2004; 68:422–428.

2 Stévant I, Kühne F, Greenfield A, Chaboissier M-C, Dermitzakis ET, Nef S. Single-cell transcriptomics of the mouse gonadal soma reveals the establishment of sexual dimorphism in distinct cell lineages. bioRxiv 2018.

3 Evans EP, Ford CE, Lyon MF. Direct evidence of the capacity of the XY germ cell in the mouse to become an oocyte. Nature 1977; 267:430–431.

4 Li L, Dong J, Yan L et al. Single-Cell RNA-Seq Analysis Maps Development of Human Germline Cells and Gonadal Niche Interactions. Cell stem cell 2017; 20:891–892.

5 Guo F, Yan L, Guo H et al. The Transcriptome and DNA Methylome Landscapes of Human Primordial Germ Cells. Cell 2015; 161:1437–1452.

6 Mayère C, Neirijnck Y, Sararols P et al. Single-cell transcriptomic reveals temporal dynamics of critical regulators of germ cell fate during mouse sex determination. bioRxiv 2019.

7 Magnusdottir E, Dietmann S, Murakami K et al. A tripartite transcription factor network regulates primordial germ cell specification in mice. Nature cell biology 2013; 15:905–915.

8 Butler A, Hoffman P, Smibert P, Papalexi E, Satija R. Integrating single-cell transcriptomic data across different conditions, technologies, and species. Nature biotechnology 2018; 36:411–420.

9 Leland McInnes, John Healy, Melville J. UMAP: Uniform manifold approximation and projection for dimension reduction. arXiv 2018.

10 Qiu X, Mao Q, Tang Y et al. Reversed graph embedding resolves complex single-cell trajectories. 2017; 14:979–982.

11 Mork L, Maatouk DM, McMahon JA et al. Temporal differences in granulosa cell specification in the ovary reflect distinct follicle fates in mice. Biology of reproduction 2012; 86:37.

12 Byskov AG, Hoyer PE, Yding Andersen C, Kristensen SG, Jespersen A, Mollgard K. No evidence for the presence of oogonia in the human ovary after their final clearance during the first two years of life. Human reproduction 2011; 26:2129–2139.

13 Hayashi K, Ohta H, Kurimoto K, Aramaki S, Saitou M. Reconstitution of the mouse germ cell specification pathway in culture by pluripotent stem cells. Cell 2011; 146:519–532.

14 Ohinata Y, Ohta H, Shigeta M, Yamanaka K, Wakayama T, Saitou M. A signaling principle for the specification of the germ cell lineage in mice. Cell 2009; 137:571–584.

15 Le Bouffant R, Souquet B, Duval N et al. Msx1 and Msx2 promote meiosis initiation. Development 2011; 138:5393–5402.

16 Nagaoka SI, Nakaki F. ZGLP1 is a determinant for the oogenic fate in mice. Science (New York, NY) 2020; 367.

17 Jan SZ, Vormer TL. Unraveling transcriptome dynamics in human spermatogenesis. 2017; 144:3659–3673.

18 Peng B, Gu Y, Xiong Y, Zheng G, He Z. Microarray-assisted pathway analysis identifies MT1X & NFkappaB as mediators of TCRP1-associated resistance to cisplatin in oral squamous cell carcinoma. PloS one 2012; 7:e51413.

19 Amoushahi M, Sunde L, Lykke-Hartmann K. The pivotal roles of the NOD-like receptors with a PYD domain, NLRPs, in oocytes and early embryo developmentdagger. Biology of reproduction 2019; 101:284–296.

20 Kissel H, Timokhina I, Hardy MP et al. Point mutation in kit receptor tyrosine kinase reveals essential roles for kit signaling in spermatogenesis and oogenesis without affecting other kit responses. The EMBO journal 2000; 19:1312–1326.

21 Chen Y, Lyu R, Rong B, Zheng Y. Refined spatial temporal epigenomic profiling reveals intrinsic connection between PRDM9-mediated H3K4me3 and the fate of double-stranded breaks. Cell research 2020; 30:256–268.

22 Findlay JK, Hutt KJ, Hickey M, Anderson RA. How Is the Number of Primordial Follicles in the Ovarian Reserve Established? Biology of reproduction 2015; 93:111.

23 Chassot AA, Bradford ST, Auguste A et al. WNT4 and RSPO1 together are required for cell proliferation in the early mouse gonad. Development 2012; 139:4461–4472.

24 Chassot AA, Gregoire EP, Lavery R et al. RSPO1/beta-catenin signaling pathway regulates oogonia differentiation and entry into meiosis in the mouse fetal ovary. PloS one 2011; 6:e25641.

25 Perander M, Al-Mahdi R, Jensen TC et al. Regulation of atypical MAP kinases ERK3 and ERK4 by the phosphatase DUSP2. Scientific reports 2017; 7:43471.

26 Pajuelo Reguera D, Cunatova K, Vrbacky M et al. Cytochrome c Oxidase Subunit 4 Isoform Exchange Results in Modulation of Oxygen Affinity. Cells 2020; 9.

27 Metzler MA, Raja S, Elliott KH et al. RDH10-mediated retinol metabolism and RARalpha-mediated retinoic acid signaling are required for submandibular salivary gland initiation. Development 2018; 145.

28 Liu C, Peng J, Matzuk MM, Yao HH. Lineage specification of ovarian theca cells requires multicellular interactions via oocyte and granulosa cells. Nature communications 2015; 6:6934.

29 Vernet N, Mark M, Condrea D et al. Meiosis initiates in the fetal ovary of mice lacking all retinoic acid receptor isotypes. bioRxiv 2019.

30 Zhao Z-H, Ma J-Y, Meng T-G et al. Single cell RNA sequencing reveals the landscape of early female germ cell development. bioRxiv 2020:2020.2005.2009.085845.

31 Paiano J, Wu W, Yamada S et al. ATM and PRDM9 regulate SPO11-bound recombination intermediates during meiosis. Nature communications 2020; 11:857.

32 Zhao F, Franco HL, Rodriguez KF et al. Elimination of the male reproductive tract in the female embryo is promoted by COUP-TFII in mice. Science (New York, NY) 2017; 357:717–720.

33 Sissaoui S, Yu J, Yan A et al. Genomic Characterization of Endothelial Enhancers Reveals a Multifunctional Role for NR2F2 in Regulation of Arteriovenous Gene Expression. Circulation research 2020; 126:875–888.

34 Yu G, Wang LG, Han Y, He QY. clusterProfiler: an R package for comparing biological themes among gene clusters. Omics : a journal of integrative biology 2012; 16:284–287.

